# Skilled lipreaders read and listen to lips

**DOI:** 10.1101/233676

**Authors:** S. Saalasti, J. Alho, J.M. Lahnakoski, M. Bacha-Trams, E. Glerean, I.P Jääskeläinen, U. Hasson, M. Sams

## Abstract

Only a few of us are skilled lipreaders while most struggle at the task. To illuminate the poorly understood neural substrate of this variability, we estimated the similarity of brain activity during lipreading, listening, and reading of the same 8-min narrative with subjects whose lipreading skill varied extensively. The similarity of brain activity was estimated by voxel-wise comparison of the BOLD signal time courses. Inter-subject correlation of the time courses revealed that lipreading and listening are supported by the same brain areas in temporal, parietal and frontal cortices, precuneus and cerebellum. However, lipreading activated only a small part of the neural network that is active during listening/reading the narrative, demonstrating that neural processing during lipreading vs. listening/reading differs substantially. Importantly, *skilled* lipreading was specifically associated with bilateral activity in the superior and middle temporal cortex, which also encode auditory speech. Our novel results both confirm previous results from few previous studies using isolated speech segments as stimuli but also extend in an important way understanding of neural mechanisms of lipreading.

## INTRODUCTION

Seeing articulatory movements on a speaker’s face amends speech comprehensible when auditory speech is hard to understand, e.g. due to acoustic noise^1–3^. Good lipreading skill is also beneficial for auditory speech processing after cochlear implantation^4^, and may provide a vital means for communication in deaf persons. To some extent, everyone can extract phonetic information from a speaker’s lips, tongue, jaw, teeth, eye brows, cheeks, and neck^5–7^, and use this ability, known as lipreading or speechreading, to support speech comprehension^8^. Only a small proportion of people become proficient lipreaders and large inter-individual variability is characteristic for lipreading skill in normal hearing adults, with most individuals falling in the lower end of the spectrum^5,9,10^. Lipreading accuracy of 45% means that an individual is five standard deviations above the mean, highlighting the difficulty of this skill^10^, when measured as a proportion of correctly recognized words from sentences by solely seeing the face of the speaker. Typically, lipreading proficiency has been gained by experience, e.g. by growing up in a hearing impaired family^8,9^. Lipreading training is an integral part of intervention when compensating for difficulties caused by mild to moderate hearing impairment, and lipreading is beneficial for supplementing hearing after cochlear implantation^11^. However, some individuals do not acquire a good lipreading skill even by training^12,13^. Better understanding of neural mechanisms underlying skilled lipreading is crucial to find better means to help such individuals.

There is evidence that differences in both low-level perceptual processing and in high-level cognitive skills contribute to variation in the lipreading skill. Well-functioning low-level processing consists of sensitive recognition of visual equivalents of phonemes (“visemes”) and sound structures in words (phonological processing). This enables better bottom-up processing of visual speech and results in more efficient lexical access^13–15^. However, low-level speech perception skills explain lipreading only partly, and high-level cognitive skills such as working memory^12^, large vocabulary^16,17^ and good inference-making^9,17^ also contribute to fluent lipreading of continuous speech. Furthermore, lipreading skill has also been found to correlate with reading ability in children^15^, but in adults only in deaf and dyslexic readers^18^. However, there are no studies of the neural basis of these behavioral findings.

Present understanding of the neural mechanisms of lipreading is based on studies using syllables, words, or single isolated sentences as stimuli. Such studies have shown that lipreading first activates the occipital visual cortical and inferior temporal areas^13,19,20^. It also activates multimodal areas in the posterior superior temporal gyrus and sulcus (pSTG/S)^21,22^ and auditory cortex including primary areas^21,23,24^ (for contradictory evidence^13,25^), as well as prefrontal speech motor areas in middle frontal gyrus (MFG), inferior frontal gyrus (IFG), supramarginal gyrus (SMG) as well as premotor cortex^20,26–29^. Thus, when we lipread, cortical areas supporting visual perception, but also those underlying auditory speech perception and speech production are employed, predominantly in the left hemisphere^28,30^. None of the above studies examined how individual differences in lipreading skill would influence brain activity.

Research on neural underpinnings of inter-individual variations in lipreading skills in hearing individuals is sparse. An early fMRI study, based on data from nine subjects, suggested that poor lipreaders had less activation in STG and MTG than good lipreaders^31^. In a more comprehensive study, 33 normally hearing subjects with lipreading skills ranging from 7% to 89% (quantified with a sentence-based lipreading test) watched silent, naturally spoken, isolated sentences during fMRI^32^. Mean activation across subjects, irrespective of lipreading skill, was found in an extensive set of areas: IFG, MFG, and the inferior parietal lobule (IPL), particularly in the left hemisphere and the MTG (peak activity in the posterior part) bilaterally. The correlation analysis suggested that small voxel clusters (4–11 voxels) in mSFG, IFG, fusiform gyrus, posterior cingulate cortex and lingual gyrus (bilaterally) were associated with lipreading skill. Using individual-thresholded activation maps and restricting the analysis to STG (outermost anatomical boundaries of Heschl's gyrus and planum temporale), the authors found statistically significant lipreading-skill related activation in the auditory cortex. This suggests that phonological processing mechanisms have a role in skillful lipreading, because STG is involved in encoding acoustic features at phonetic level^33–38^.

In a similar vein, Capek et al.^39^ found that activation in the left posterior STG as well as MTG and MFG was positively associated with the lipreading skill in deaf individuals but not in normal-hearing subjects. In the latter, they found lipreading-skill dependent activity in the right lingual, posterior cingulate, right postcentral and inferior temporal gyri, suggesting superior articulatory and face processing skill. Notably, the deaf subjects were better lipreaders than normal hearing subjects, which probably explains the results. The stimuli were single words that require less linguistic processing than sentences^32^. The linguistic complexity might affect the strength of cortical activity, as lipreading of sublexical structures such as pseudowords and syllables has been found to activate areas sensitive to visual motion more than lipreading words and sentences, which requires more lexical lipreading^25^.

In real life, lipreaders need to comprehend continuous natural speech, which requires meaning-based processing of language and integration of linguistic information over extended time periods. To imitate real-life speech, we used here for the first time a natural complex narrative as a stimulus. Comprehension of the narrative requires efficient joint utilization of information from simple phonetic features up to those necessary to integrate information over longer time scales. Our aim was to 1) reveal brain areas where activity is predicted by the lipreading skill, and 2) to characterize to what extent lipreading engages similar neural mechanisms in a similar manner as listening and reading the same narrative. To this end, subjects with large inter-individual variation in lipreading skills had their brain activity measured with 3-T fMRI while they 1) lipread the eight-minute narrative from a silent video showing the speaker’s face, 2) listened to the same narrative without seeing the face of the speaker, and 3) read a time-locked transcript of the narrative (**Fig. 1**). We used intersubject correlation (ISC) of the blood oxygenation level dependent (BOLD) signal time courses to identify similarity of brain activity between different narrative types (**Fig. 1**). ISC is a data driven method suitable for studying brain activity triggered by naturalistic, complex stimuli such as long-duration movies and narratives^40–43^. It is based on voxel-wise correlation between the BOLD signal time series of the subjects. We used a between-condition ISC to test similarity of brain activity in the three narrative conditions: BOLD signal time courses from each condition (lipreading, listening, reading) were used as a model to identify brain areas with similar time courses in another condition.

**Figure 1.**
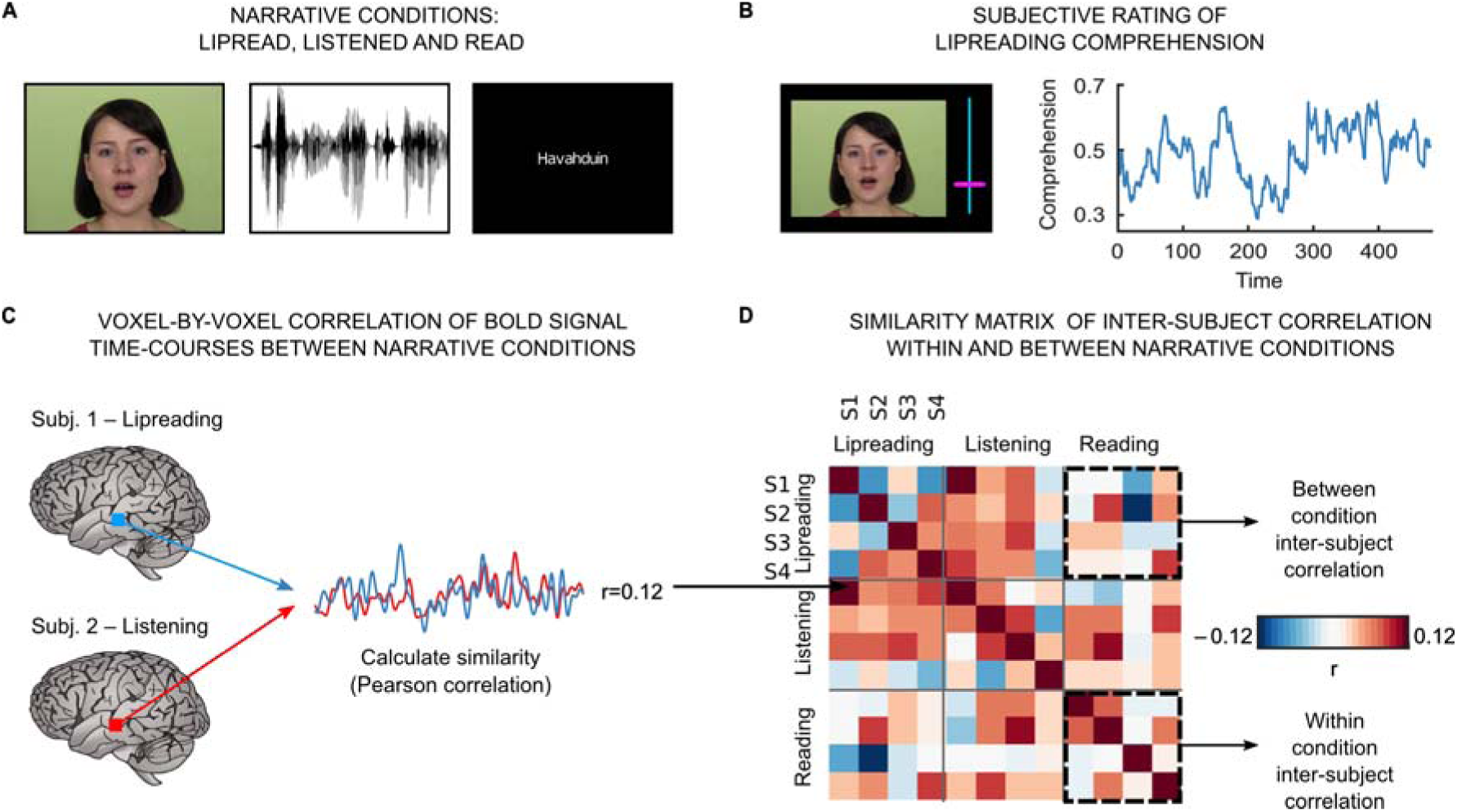
Illustration of the experimental design and between-narrative-type intersubject correlation. A) The subjects lipread, listened to, and read the same narrative. B) After scanning, the subjects lipread the narrative again and rated how well they comprehended what the speaker said. C) We used BOLD signal time courses of Subject 1 during Listening as a model to identify brain areas with similar time courses of Subject 2 during Lipreading: we computed the *r* statistics voxel-by-voxel between the signals and repeated the process for all subject pairs. D) Similarity matrix of four subjects depicting between- and within-condition intersubject correlations from one voxel in the posterior temporal cortex. In contrast to previous studies, we analyzed the between-condition ISCs.

We hypothesized that inter-individual variability in lipreading skills would depend on differential use of the same neural mechanisms in the temporal cortex, which are also used in coding heard speech. More specifically, if skilled lipreading would depend on superior low-level linguistic abilities such as phonological processing, good lipreaders would have similar brain activity in superior temporal areas and/or inferior-parietal and inferior-frontal areas^32,33,44^. Alternatively, if good lipreading skill depends more on higher-level cognitive skills, such as efficient semantic processing and inference making^9,12,17^, activity would be more similar in middle temporal and/or fronto-parietal areas. We also hypothesized that lipreading continuous speech would be largely supported by the same neural mechanisms that underlie linguistic processing during listening and reading. Additionally, we expected to find neural activity during lipreading that is revealed only using natural narrative.

## RESULTS

### Lipreading skill of subjects

To ensure a high variability in the lip-reading skills, subjects were pre-screened with a web-based lipreading test. Then, prior to the fMRI scanning, subject's lipreading skills were estimated with a previously validated test, which consisted of 10 sentences of variable length (Lonka, 1993). Inter-individual variability was very large with the number of correctly recognized words ranging from 3 to 50 out of 50 words (mean = 25; SD = 13). After the fMRI scan, subjects used a dynamic rating tool (see METHODS) to provide a continuous subjective estimate of how well they could lipread the stimulus narrative. The estimate varied over time (**Fig. 1B**). The mean, calculated over the whole narrative, ranged from 0.016 to 0.98 (scale 0–1). Linear regression showed a significant (r = 0.54, p < 0.003) correlation between lipreading score and mean subjective rating of comprehension (**Fig. 2**), demonstrating that lipreading score predicted well the comprehension of the narrative in the scanner.

**Figure 2.**
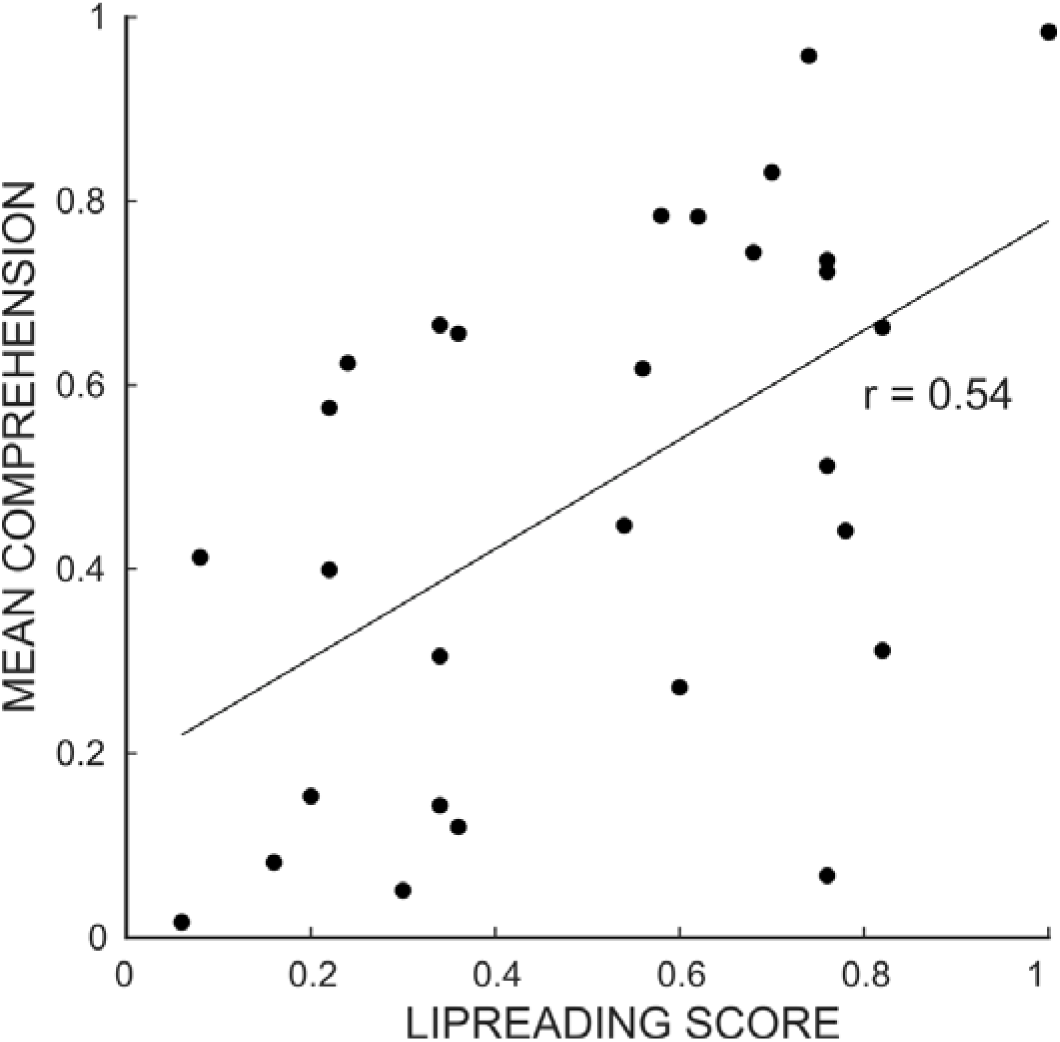
Correlation of two different behavioral measures of lipreading skill. Performance in a sentence-based lipreading test predicted well the subjective comprehension of lipreading the long narrative.

### Brain imaging results

#### Similarity of brain activity during listening, reading and lipreading

To disclose the similarity of brain activity during lipreading and the other two conditions, we measured between-condition ISC of BOLD signals. In the analysis, brain activity during lipreading functioned as a model to identify brain areas with similar BOLD signal time courses between listening and reading. We also calculated the similarity of brain activity during reading and listening. Data were cluster-corrected with FSL randomize with 5000 permutations (**Fig. 1**, see METHODS for details).

The brain activity during lipreading and listening (**Fig. 3A**, See also Table S1 for peak values) was significantly similar bilaterally in the middle and posterior superior temporal gyrus and sulcus (STG/S), as well as along the whole bilateral middle temporal gyrus and sulcus (MTG/S) extending to the temporal pole (TP), left superior marginal gyrus (SMG), right inferior frontal gyrus (IFG), and in two areas of the left primary motor cortex (M1) corresponding approximately to the hand^45^and mouth representation areas^46^. Similar activity was also observed in bilateral precuneus (PCUN), occipital midline areas around the calcarine sulcus (CS) (extending to the lingual gyrus (LG) and the right fusiform gyrus (FG)), right-lateralized areas of cerebellum and left somatosensory cortex.

**Figure 3.**
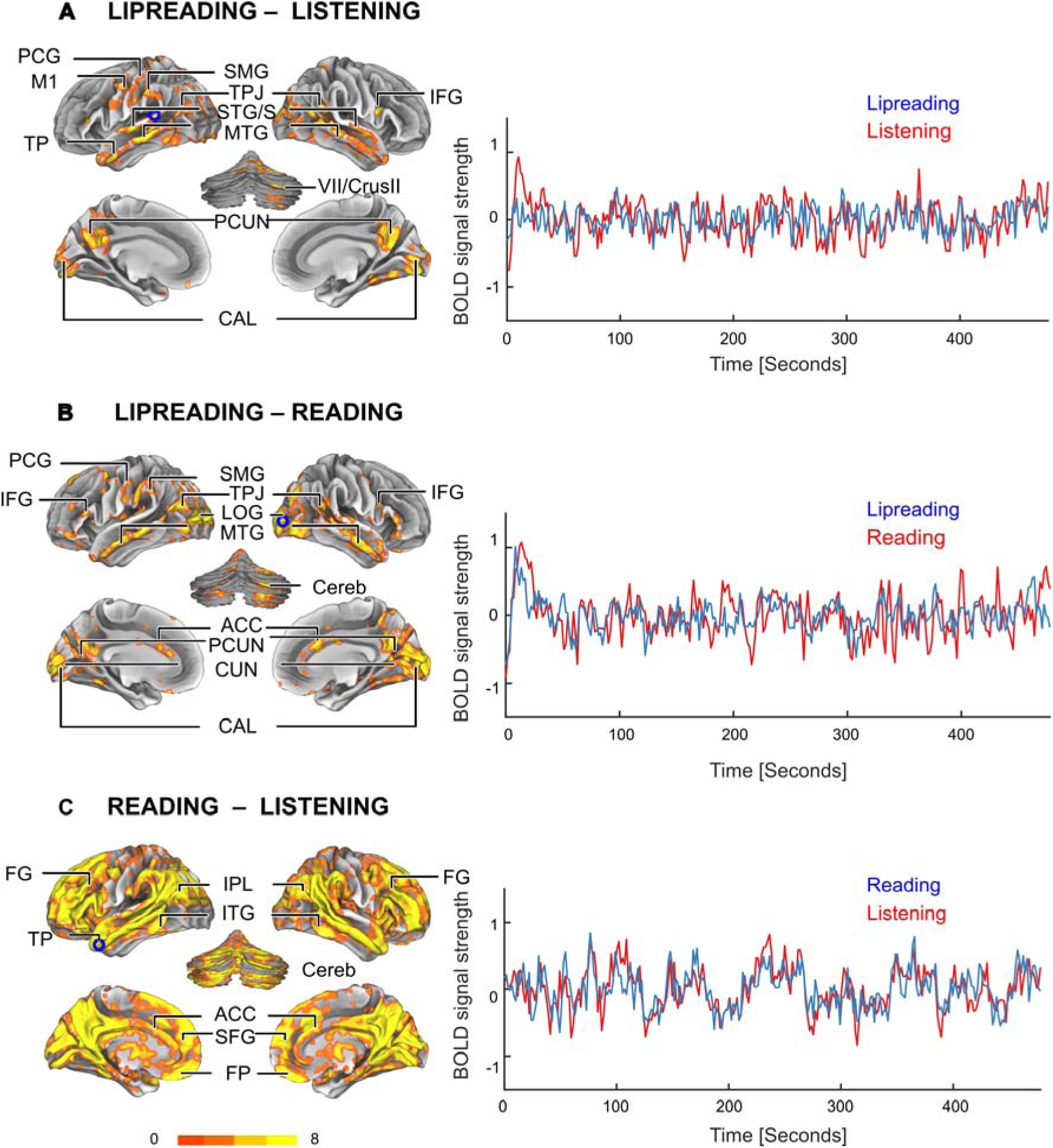
Inter-subject correlation between narrative types. Left: Brain areas showing similar activity during processing of two narrative types (permutation-based cluster-correction, p_*corrected*_ < 0.05). Right: Time series of peak values of BOLD signal strengths from the largest cluster (denoted with a blue circle) where the signals were significantly similar. A) During lipreading and listening, similarity was largest in the left-hemispheric temporal cortex. Maximum similarity (denoted with a circle) was centered at MNI coordinates x = -62, y = -22, z = - 4. B) During lipreading and reading, similarity was largest in bilateral visual-cortical areas. Maximum similarity was centered at MNI coordinates x = 22, y = - 98, z = 12 C) During listening and reading, BOLD signals were similar in large parts of the brain. Maximum similarity was centered at MNI coordinates x = -54, y = 10, z = -30.

During lipreading and reading, brain activity was significantly similar (**Fig. 3B**, see also **Fig. S1**) along the whole MTG/S extending to TP, left SMG, bilateral lateral occipital gyri (LOG), starting from the occipital pole (OP), and extending around CS, comprising primary visual cortex as well as other early visual areas. Furthermore, brain activity was similar in left and right cerebellum, PCUN and cuneus as well as in anterior cingulate (ACC).

During reading and listening, brain activity was significantly similar in an extensive set of brain areas *excluding* primary somatosensory cortex, ventrolateral visual cortex, subcortical areas (not shown in the figure), frontal orbital cortex, and right supra-temporal auditory cortex (**Fig. 3C**; see also **Fig. S1**). In contrast to lipreading, the content of the message was comprehended equally accurately by both listening and reading, which probably explains the large-scale similarity in brain activity. Due to the poor signal-to-noise ratio and inter-individual variability of BOLD signals during lipreading and variability in the time courses of comprehension ratings and their relative unreliability at a single-subject level, a more "direct" analysis where brain activity during lipreading would have been predicted by experienced comprehension, was not feasible.

To illustrate brain areas that had similar activity during all narrative types, **Fig. 4** depicts the overlap of the three between-condition ISCs (**Fig. 3**). These areas comprise the bilateral anterior MTG and left middle MTG, posterior STS, bilateral IFG, left temporo-parietal junction (TPJ), bilateral CS, bilateral PCUN, bilateral orbitofrontal cortex (OFC) as well as the left superior frontal gyrus (SFG) and right cerebellum (VI and VIIA/Crus II).

**Figure 4.**
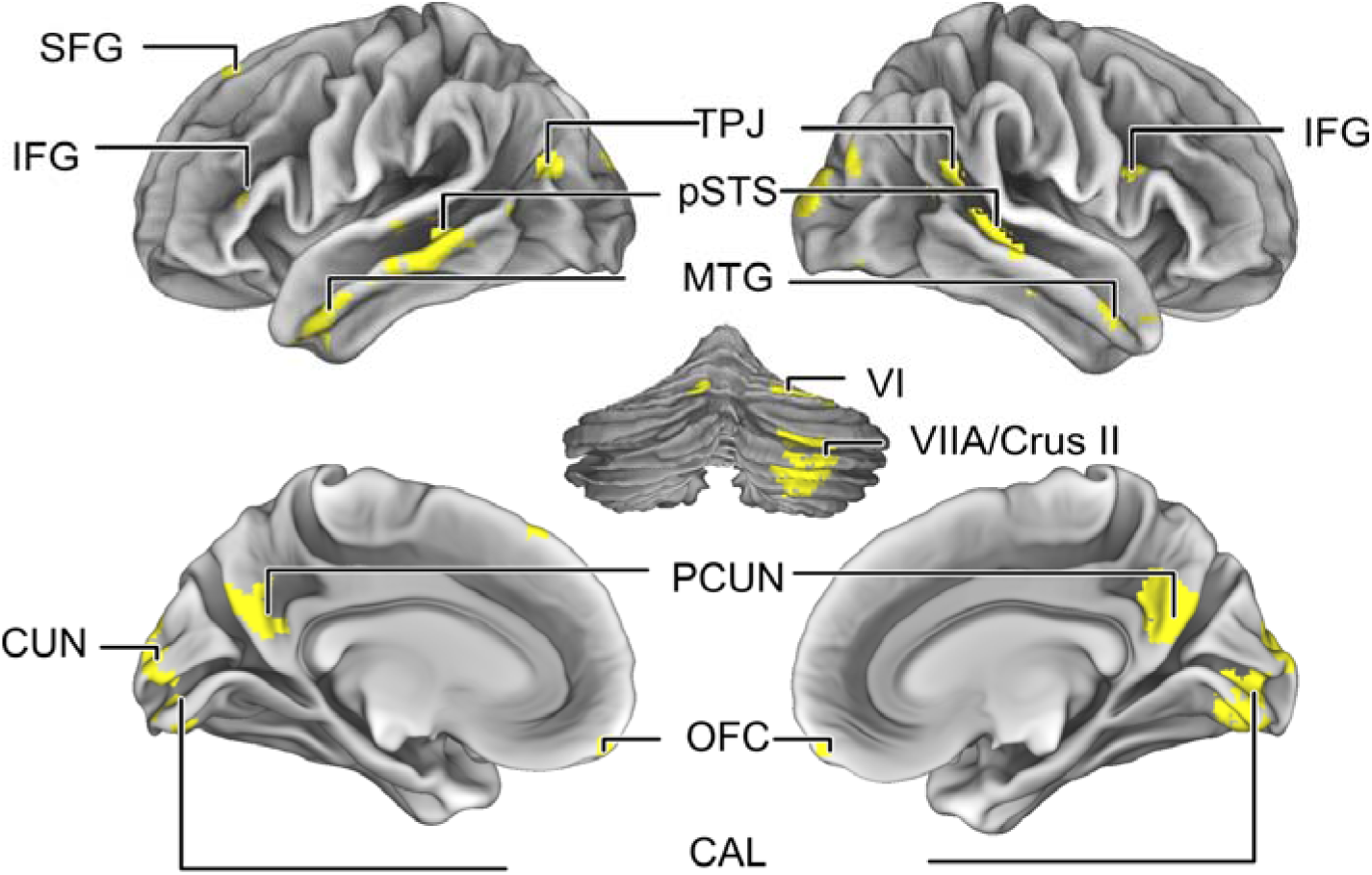
Brain areas showing overlap for all between-condition ISCs: Similarity was observed in the left STG, and left SFG. Areas showing similarity bilaterally were in the posterior STS and anterior MTG, IFG, and PCUN, as well as in right cerebellum (VI and VIIA/Crus II).

#### Lipreading skill predicting brain activity

Lipreading skills of our subjects varied extensively, ranging from very poor to excellent (**Fig.2**). Because lipreading scores correlated strongly with the subjectively experienced comprehension of the narratives, we used the latter to identify brain areas, where similarity of activity was dependent on the lipreading skills. We did this by estimating how well experienced comprehension predicted the similarity of brain activity during lipreading and listening (see **Fig. 3a**).

ISC between lipreading and listening was linearly regressed with the mean behavioral subjective rating of lipreading comprehension and data were thresholded with permutation-based cluster-correction (p_*corrected*_ < 0.05). Similarity of brain activity during lipreading and listening was dependent on lipreading performance bilaterally in middle STG/S and MTG and in the anterior STS and posterior MTG in the right hemisphere (**Fig. 5**). These temporal-cortical areas are typically active when listening to speech. Similarity did not extend to the primary auditory cortex (see also **Fig. S1**).

**Figure 5.**
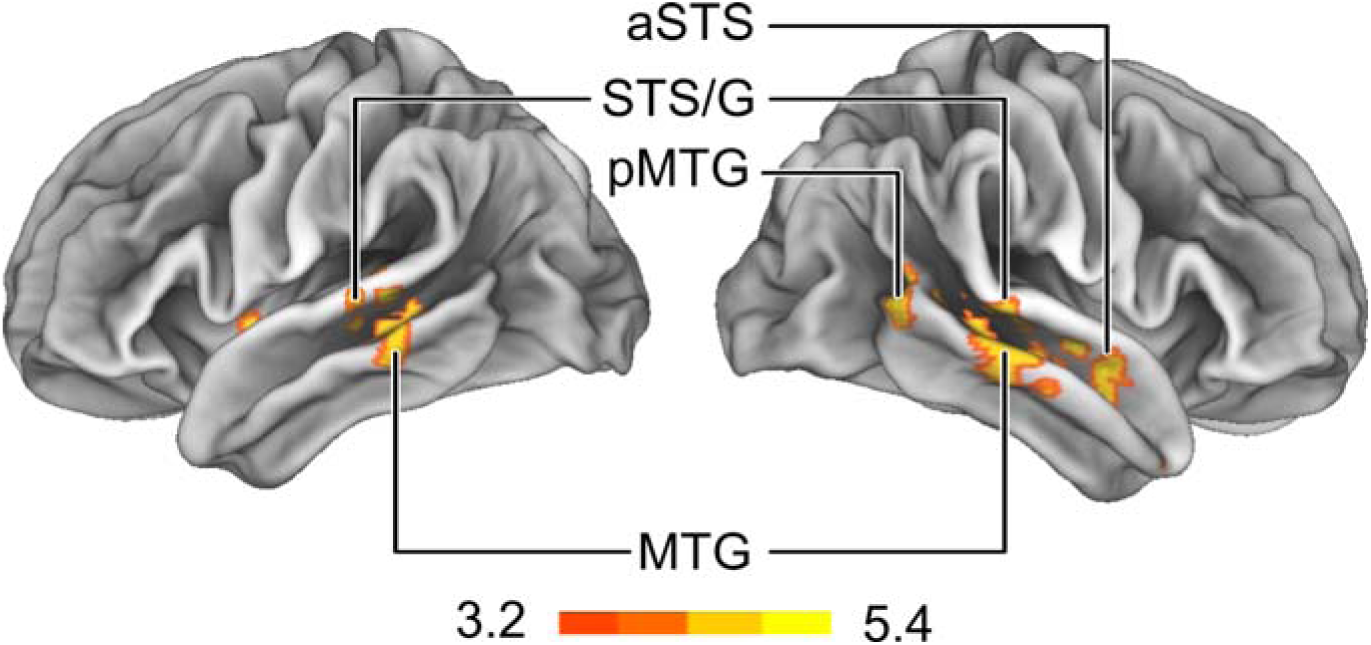
Brain areas showing higher similarity of activity during lipreading and listening in subjects with higher lipreading skills. Higher pairwise subjective rating of lipreading comprehension predicted similar brain activity bilaterally in middle STG/S and MTG as well as posterior MTG and anterior STS in the right hemisphere (permutation-based cluster-correction p_*corrected*_ < 0.05).

## DISCUSSION

The current study pioneers in studying neural mechanisms of lipreading rich natural narrative in subjects with large inter-individual variability in lipreading skills. We demonstrate that skilled lipreaders are characterized by recruitment of the same temporal-cortical brain areas that support phonological processing when listening to speech in the left hemisphere. Importantly, analogous and even more extensive recruitment was found in the right hemisphere. In relation to other narrative types, we found that lipreading activated a set of brain areas that were also activated during listening and reading the same narrative. These areas are in the anterior MTG, IFG, posterior STS, TPJ, and PCUN bilaterally and around Crus II in the right cerebellum. In addition, lipreading activated auditory areas predominantly in the left STG similarly to listening to the narrative, and midline visual-cortical areas similar to reading the narrative.

Our new results confirm the previous finding on the significant relationship of left STG activation during lipreading isolated sentences and performance in the lipreading test^31,32,39^. In addition, lipreading skill predicted activity in the auditory areas in right STG, bilateral middle STS and MTG, as well as in right posterior MTG and anterior STS. Previous studies have identified STG as the core cortical region for processing acoustic features of speech and in assessing sublexical structures such as phonetic characteristics of words and syllables^35,37,47^. High-density direct cortical surface recordings during listening to continuous auditory speech have shown that the STG contains neurons selectively responding to phonetic features^37^. Importantly, our results show that when lipreading a rich natural narrative, activity in posterior MTG — which has been implicated in processing of meaning^33,44,48^ and in lexical-syntactic information retrieval^49,50^ — becomes visible. Comprehension of natural narrative requires constantly integrating new semantic information with earlier context. Therefore, our novel finding of lip-reading-dependent activity in MTG activity likely reflects subsequent selection and integration of semantic information, following initial lexical access.

Importantly, the lipreading-skill-dependent temporal-cortex activity was bilateral, and even more pronounced in the right hemisphere. Naturalistic auditory speech stimuli such as narratives constantly elicit more bilateral brain activity than simpler stimuli^51–54^, and now we show that this is also true for lipreading. Campbell et al.^55^ speculated that a good lipreader may recruit neural mechanisms in the right hemisphere, but previous studies have not provided any experimental support for this. The right temporal cortex has been suggested to be specialized in processing intonation-level time intervals, therefore perception of prosodic features of speech involves the right hemisphere^34,35,47,56^. Accordingly, lipreading-skill-related activity in the right STG/S and MTG could be at least partly related to tracking visible prosodic features such as head and eye-brow movements^57,58^.

Our data support the view that neural mechanisms decoding low-level auditory speech features (phonological processing) contribute to skilled lipreading. Related activity in areas sensitive to semantic processing such as lexical access is likely enabled by these low-level mechanisms. By contrast, our data do not suggest that higher-order cognitive functions would be the key mechanisms for skilled lipreading, because the skill-dependent brain activity was localized to restricted areas in bilateral temporal cortex.

When lipreading skill was not taken into account, we found lipreading-related activation not only in the multimodal posterior STS and STG^22,30^, but it extended to more anterior and lateral auditory cortical areas in the STG and MTG that support phonetic processing^35–38^. Anterior parts of the STG and MTG have been implicated in mapping acoustic phonetic cues into lexical representations^59,60^. Thus, the current results suggest that same mechanisms that are involved in sound-based coding of phonemes in mid-STG and MTG respond also to visual speech gestures. Importantly, the similarity of activation between lipreading and reading did not involve the STG, suggesting that phoneme analysis is not similarly needed in reading text as it is in lipreading. However, subjects of the current study were all adult individuals with fluent, automated reading skills, whereas the association of lipreading and reading was found in hearing impaired and deaf children^18,61^. It is quite possible that the role of phonetic processing diminishes during intact learning to read.

Previous research using simple stimuli such as syllables and words has suggested that lipreading activates primary auditory cortex in the left hemisphere^23,24,30^. We did not find such activity here (Fig. S1). In the present study, responses to auditory and visual transients, such as naturally occurring pauses in the narrative, were regressed out from the BOLD signals, as they are known to trigger activity in widespread cortical areas and in the primary sensory areas^62^. It is quite possible that auditory/visual (?) transients, but not linguistic stimulus features, caused primary auditory-cortical activation in the above-mentioned studies (see also^5^).

Our data rather suggests an extensively shared mechanism for lipreading and listening to speech, and does not support a suggestion that lipreading mainly relies on intact mechanisms subserving visual perception^5,63^ or that during lipreading, a specific visuomotor pathway, involving the middle part of the left MTG and frontoparietal motor areas, mediates speech motor control^64^.

Previous studies with isolated simple linguistic stimuli have shown that brain areas related to speech production are activated during lipreading^27–29,65^, which is also supported by our data. Premotor cortex and posterior STG/S have been suggested to form a network that track visual speech features. In line with that, we found that lipreading activated the primary motor cortex in the left hemisphere similarly to listening (**Fig. 3A**), but not reading (**Fig. 3B**). Apparently, motor knowledge of articulatory gestures modulates auditory-cortical processing through reciprocal sensory-motor connections^20,28,65,66^ and motor knowledge of our own speech production is used in lipreading others^25–27,30 64^.Furthermore, activity in inferior parietal regions around the temporo-parietal junction (TPJ; **Fig. 3A, 3B**, and **3C**) could be related to accessing the stored motor representations of speech during visual speech gesture decoding^21^.

Temporo-parietal brain areas that showed similar activity for all three narrative types have previously been identified as an integrated system underlying high-level language processing^53,67–70^. Activity in the MTG has previously been shown to be related to meaning-based processing (semantics)^47,71^ and here, the similarity of activity in all conditions suggests that these areas supported comprehension of the meaning in all narrative conditions. This is in line with previous studies that have suggested similar responses to listened and read narratives^72^ as well as to listening to the same narrative in different languages^52^. The similarity of activity in the bilateral posterior STG and PCUN, as well as in the left IFG may reflect processing of complex language (grammar) and context^51,69^ as well as mentalizing^73–76^. Also, the activity of right cerebellum (VI and VII/Crus II) during all narrative types confirms previous findings that it is not only involved in sensorimotor tasks, but also in complex linguistic processing^77–80^.

Similarity of brain activity during listening and reading was much more extensive than that between lipreading and listening or lipreading and reading. Similarity extended to the frontal and midline structures as well as to the bilateral cerebellum (**Fig. 3C**). Similarity was lacking or was clearly smaller in the right auditory cortex in STG, visual cortices as well as somatomotor areas. Even when the complex and equally-well understood linguistic contents is received *via* different sensory modalities, its neural processing is largely similar, also in the left auditory-cortical areas, excluding the primary auditory cortex (see **Fig. S1**). Involvement of such a high number of brain areas suggests a comprehensive and similar recruitment of different cognitive operations, as e.g. experiencing emotions, imagery related to the contents of the story, social reasoning as well as episodic and self-referential memory^69,81–83^.

The current study is the first to study lipreading using a natural long-duration narrative, which requires processing of many types of information in the context of the storyline. As is evident in the activity depicted in **Fig. 3C**, if such a stimulus is unambiguous, it activates most of the brain bilaterally including large areas of cerebellum, the exact activation pattern depending on the context of the narrative^75,76^. Lipreading is a much more difficult task than listening to speech^6,10^, due to the similarity of lip shapes of different sounds (e.g. bilabials /m/ and /p/) and the poor visibility of some other sounds (e.g. velopharyngeal sounds /k/ and /h/ that are produced inside the oral cavity). Only a subset of subjects in the present study reported that they comprehended the narrative well. This is paralleled by the much more restricted activation of the brain during lipreading than during listening to speech or reading (see also^65^).

The presentation sequence of three narrative types was counterbalanced across the subjects, as described in Methods. However, because of our use of within-subject design, for example previous listening to the narrative might have influenced neural processing during reading or lipreading the narrative. This might have, due to adaptation and familiarity, weakened the ISC between listening and reading which, however, was extensive. In addition, reading or listening to the narrative before lipreading might have made lipreading easier for some subjects and modified corresponding neural processing. This could have somewhat influenced the results of our analysis on the effect of lipreading skill on ISC. However, our use of subjective rating of the lipreading difficulty after scanning probably mitigates such effects.

Lipreading the narrative in the present study was a difficult task probably requiring the subjects to use their individual knowledge and strategies in interpreting the message. Such idiosyncrasy obviously leads to inter-individual differences in neural processing and low between-subject similarity of brain activity. Between-condition ISC, however, allowed us identify similarities in brain mechanisms related to lipreading. To address the differences of inter-subject correlation between skilled and poor lipreaders more directly, further studies should use a large group of excellent lipreaders, or alternatively could study a set of good lipreaders several times to calculate intra-subject correlations. The latter approach could illuminate the possible idiosyncratic strategies of skilled lipreaders.

As a conclusion, our results suggest that after initial visual processing, lipreading is supported by a set of same brain areas involved when listening to speech. Skilled lipreading is associated with activity in the bilateral auditory temporal cortex, suggesting an efficient coding of visual speech gestures by the same mechanisms used in auditory coding of phonetic speech features. Listening and reading a natural narrative activate the brain extensively and similarly. However, similarity of brain activity during lipreading vs. reading or listening the same narrative is much less extensive.

## METHODS

### Subjects

The volunteer subjects were 31 healthy native Finnish-speaking females (mean age 30.9 years, range 20–49). The data of one subject was removed due to excessive head movement and of another one due to poor attention (eyes closed during scanning for approximately three minutes), resulting in a final sample of 29 subjects. All subjects were right-handed (Edinburgh handedness inventory^84^) and reported normal hearing and normal or corrected to normal (with contact lenses) vision, and no psychiatric or neurological disabilities. All subjects signed an informed consent, and received monetary compensation for their time. The study was approved by the research ethics committee of Aalto University and it was conducted in accordance with the Helsinki Declaration for Human studies.

We developed an online test for screening lipreading skill. We first recorded a female speaker with clear visual articulation, speaking 100 sentences that were translated to Finnish from the *CID everyday sentences* with varying length and sentence structure^85^. Out of 100 sentences 10 sentences that mimic the sentence structure of Finnish language well, were chosen for an online screening tool that was distributed via student mailing lists (speech therapy and sign language interpreters), and via the *Federation for Hard of Hearing in Finland*. The subjects were instructed to type down the words they recognized from the silent videos. The number of correctly recognized words out of the total of 57 words was used as each subject's lipreading skill score. For the current study we then chose individuals with a large inter-individual variation in their lipreading skills, and their lipreading was further confirmed on site with another lipreading test^86^.

### Stimuli and experimental design

The stimulus was a narrative (duration 7 min 54 seconds) told by a female speaker, portraying the events and thoughts during her day from a first-person perspective. The control stimulus was an unintelligible, gibberish version of the same narrative, which was created by replacing consonants from each word of the original narrative with other consonants with similar place of articulation, but the suffixes that indicated syntax were kept unchanged. This resulted in a meaningless string of speech sounds that had very similar acoustic properties and structure (syntax) than the original narrative, but no content (semantics), sounding phonetically natural. Results related to the gibberish narrative will be reported separately. The speaker, who was chosen for her clear visual articulatory gestures, was video-recorded reading the narrative aloud from a prompter in an acoustically shielded room using an additional Sennheiser EW 112P-G3-C - microphone and Canon XA10 video camera. The speaker had rehearsed reading the stories aloud at home. Two external LED lights (Dyna-Core Elf2-DS LED) illuminated the speaker’s face in front of a pale green background canvas to provide good visibility of articulatory movements.

The stimuli were edited with Matlab (MathWorks Inc.). Each paragraphs starting points were matched by stretching the audio waveform to maximize the similarity of the loudness envelopes. Playback speeds were between 95% and 101% of original speed during the paragraphs to keep the changes in playback speed unnoticeable to the subjects (higher deviations from normal playback speeds were allowed during pauses between paragraphs). The root mean square (RMS) envelopes of the two stimuli after this transformation were highly correlated (r=0.57 for RMS in 0.5-second windows; r=0.88 after convolution with a canonical hemodynamic response function (HRF)). The video recording was separated into visual and audio files. Silent videos of the speaker’s face on a light green background were used in lipreading condition. In auditory condition, narrative was presented with a blank screen in a similar shade of green as in the background of the lipreading condition.

We also created a written a 614-word transcript of the narrative. Written words were presented centrally on the screen time-locked to each word of the original spoken narrative. When the duration of the words in the spoken narrative were very short, two or three words were presented simultaneously to keep the timing.

Stimuli were presented using Presentation software (Neurobehavioral Systems Inc., Albany, California, USA). The audio stimuli were played with an MRI-compatible in-ear earbuds (Sensimetrics S14 insert earphones). In addition, MRI-safe protecting earmuffs were placed over the earbuds for noise removal and safety. Sound intensity was adjusted for each subject to be loud enough to be heard over the scanner noise by playing example stimuli that were normalized to the same level as the auditory stories during a dummy EPI sequence before the actual experiment. In the MRI scanner, the stimulus videos and texts were back-projected on a semitransparent screen, using a Panasonic PT-DZ110XEJ projector (Panasonic Corporation, Osaka, Japan). The viewing distance was 35 cm, the width and height of the projected face image was 380 pixels in height and 161 pixels in width corresponding to approximately 10.9° vertical and 4.6° horizontal angle in the visual field.

The stimulus sequence consisted of six different narratives, three intact used as stimuli in the present paper, and three gibberish versions of the same narratives. The latter three always preceded the corresponding intact narratives, which were then presented in an order counterbalanced across the subjects (**Fig. 1A**). Each stimulus presentation began with a fixation cross in the middle of the screen.

#### Assessment of lipreading skills

Prior to scanning, each subject’s individual lipreading skills were confirmed on site with a sentence-based lipreading test, comprising 10 sentences of variable length^86^. The test stimuli were presented on 17” computer screen (resolution 1366 × 768) with 40-cm viewing distance. The face image (height 9.6 cm width 6 cm) on the screen corresponded 12.8° vertical and 8.5° horizontal angle in the visual field. The speaker repeated each of the 10 sentences twice: first by saying it with a slower-than-usual speech rate, and then with normal speech-rate. Subjects were instructed to write down words they were able to recognize. The number of correctly recognized words (out of maximum 50) provided the lipreading skill score. For example, if the sentence was “On Thursday we eat pancakes” and a subject wrote down “we eat”, he got a score of 2 out of 5.

Immediately after the fMRI experiment, the subjects rated their comprehension of the silent visual narrative (continuous scale from very poor to very good). The instruction was to estimate how they comprehended the narrative when it was first presented to them in the scanner. This was done using a web-based dynamic rating tool (https://git.becs.aalto.fi/eglerean/dynamicannotations/tree/master, see^87^. Data were collected at 5 Hz. Subjects used a mouse to move a small cursor at the right side of the screen up (good comprehension) and down (poor comprehension). The original scale of rating was from 0 (very poor) to 1 (very good). Rating was done after fMRI acquisition to prevent any influence on the neural activity during scanning.

#### MRI acquisition

Functional magnetic resonance imaging (fMRI) was done with a 3T Magnetom Skyra whole-body scanner (Siemens Healthcare, Erlangen, Germany) and a standard 20-channel receiving head/neck coil at the Advanced Magnetic Imaging (AMI) Centre of the Aalto NeuroImaging (ANI) infrastructure at Aalto University School of Science. For functional scans, images were acquired using a T2-weighted echo planer imaging (EPI) pulse sequence: repetition time (TR), 1700 ms; echo time (TE), 24 ms; flip angle, 70°, each volume comprising 33 slices of 4 mm thickness with 0 mm gap. A total of 295 volumes were acquired from which 13 volumes were discarded from each run to exclude brain activity during the viewing of pre-stimulus fixation cross and after ending actual stimulus presentation. In-plane resolution was 3 × 3 mm^2^ (field of view (FOV), 192 × 192mm^2^). Anatomical T1-weighted structural images were acquired at a resolution of 1×1×1 mm^3^ (MPRAGE pulse sequence, TR 2530 ms, TE 3.3 ms, TI 1100 ms, flip angle 7°, 256 × 256 matrix, 176 sagittal slices).

To monitor subjects' attention, their eye gaze was recorded with an EyeLink 1000 eye tracker (SR Research, Mississauga, Ontario, Canada; sampling rate 1000 Hz, spatial accuracy 0.5°). Prior to the experiment, a nine-point calibration and validation was performed.

### Data Analysis

#### Preprocessing

The fMRI data were preprocessed with FSL software (www.fmrib.ox.ac.uk/fsl) using the BRAMILA parallel preprocessing pipeline (https://version.aalto.fi/gitlab/BML/bramila). First, after correcting for slice-timing during acquisition, the EPI volumes were spatially realigned to the middle scan by rigid body transformations to correct for head movements using FSL MCFLIRT. EPI and structural images were co-registered and normalized to each individual’s anatomical scan (linear transformation with 9 degrees of freedom with FSL FLIRT; structural images were cleared from non-brain tissues with FSL BET) followed by a linear transformation from anatomical to standard MNI template space (12 degrees of freedom; FSL FLIRT). Finally, BOLD time series were detrended (linear detrend), motion parameters were regressed out (24 parameters expansion^88^) as well as average signals at deep white matter, ventricles and cerebro-spinal fluid^88^. Finally, a temporal high-pass filter with a cut-off frequency of 0.01Hz was applied, followed by spatial smoothing with a Gaussian kernel of 8mm FWHM.

#### Similarity of brain activity between narrative types measured with inter-subject correlation (ISC) of BOLD signal time courses between narrative conditions

Our main interest was to find similarities in brain activity during processing of the same narrative presented *via* different means. The data were analyzed with voxel-wise comparison of the BOLD signal time courses triggered by the listened, read, and lipread narratives. We estimated the similarity of the time series using inter-subject correlation analysis (ISC^41^), which examines the temporal similarity of the signals in individual voxels. ISC is a data driven method suitable for more ecologically valid stimulus paradigms. It is optimal for analyzing data acquired from experiments with complex stimuli^40,42,43^. A recent paper has shown that the stimulus structure can have an effect on ISC^62^. Therefore, as recommended, we controlled the possible effect of silent pauses by modelling the stimulus structure based on the presence of speech as in Lahnakoski et al.^87^. ISC was calculated using the ISCtoolbox^42^. First, inter-subject correlation matrices were obtained for each brain voxel by calculating all pairwise Pearson's correlation coefficients (r) of the voxel time courses across the subjects, resulting in 406 unique pairwise r-values for each voxel in each condition. Here, however, we performed ISC between conditions: BOLD signal time courses from each condition (listening, reading and lipreading) were used as a model to identify brain areas with similar time courses in another condition (**Fig. 1 D**). This provided a measure of which brain areas during lipreading responded similarly with the brain responses measured during listening or reading. Specifically, given a subject pair with one subject in condition *c1* and a second subject in condition *c2*, for each voxel we compute the average *r*statistic as the average of all pairwise correlations between the BOLD signal time series *s*(t) at the voxel (formula 1). Unthresholded brain maps for all are found in Neurovault (/collections/BJRMDQXU/).

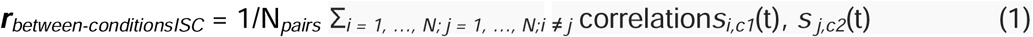

To obtain statistical significance of the between-condition ISC and to control for multiple comparisons, whole brain permutations (N=5000) were performed with FSL randomize with the threshold-free cluster enhancement option^89^.

#### Predicting brain activity with lipreading skill

To reveal the brain areas related to lipreading proficiency, we predicted the ISC between listening and lipreading against the individual subjective mean rating of comprehension of the silent narrative. For each voxel, the pairwise BOLD similarity between two subjects in the two different condition is compared to a pairwise comprehension score based on the joint lipreading comprehension (average subjective rating of comprehension between the two subjects). To test for significance of the association between similarities, we run a Mantel test^90^, which is a non-parametric test where subjects labels are shuffled at each permutation. To obtain statistical significance of the association and to jointly control for multiple comparisons, whole brain permutations (N=5000) were performed with FSL randomize with the threshold-free cluster enhancement.

## Author Contributions

SS: planned the experiment, prepared stimuli, collected and analyzed data, prepared figures, wrote the manuscript as first author.

JA: planned the experiment, collected and analyzed data, wrote the manuscript.

JL: planned the experiment, prepared and edited stimuli, analyzed data, wrote the manuscript.

MB-T: planned the experiment, wrote the manuscript

EG: developed methods, analyzed data, wrote the manuscript

IJ: planned the experiment, supervised collecting data, wrote the manuscript

UH: planned the experiment, wrote the manuscript

MS: principal investigator of the study, planned and supervised the experiment, wrote the manuscript

None of the authors claim any conflict of interest. None of the authors report any competing financial interests.

## Acknowledgments

We kindly thank the subjects for their participation in the experiment. We also thank personnel of imaging facilities in AMI centre of Aalto University School of Science, Espoo Finland, and especially nurse Marita Kattelus for her help in fMRI imaging. The current study was performed with the support of Alfred Kordelin -foundation (personal grant for the first author), Finnish Cultural Foundation (grant number: 150496 to JML) and Academy of Finland (grant number: 276643).

## Appendix A. Stimulus narrative translated to English

I woke up to the persistent sound of my alarm clock. Waking up felt heavy, but I forced myself out of bed. I turned off the nasty sound of my alarm clock and stretched for a moment. Half asleep I made my bed and pulled a blouse and trousers on me. After opening the curtains, I felt considerably more awake. Light that was rushing in told me that spring had advanced. I turned around and headed towards the bedroom door. On my way, my foot hit a sauna bucket which was lying behind the bed and made a ruckus when falling. I had to curse for a moment when the pain hit my toes, but then I lifted the bucket up and headed towards the kitchen.

I went straight to the fridge. Jar of yoghurt and pear were an adequate breakfast. For some reason, my husband Jarkko’s mobile phone was also in the refrigerator. I was beginning to be in a hurry for work, but I nonetheless brewed some coffee, the aroma of which floated delightfully into my nose. Suddenly I felt hands begin to rub my shoulders. Jarkko had appeared behind me and he managed to surprise me pleasantly. When I wondered why Jarkko had not yet left for work, he replied that he had felt nauseous during the small hours. He was going to go right back to rest.

While sipping the last of my breakfast coffee, I gave the weary Jarkko a kiss. I pulled a jacket on me, I put my shoes on and stepped outside. As the door opened, beautiful birdsong filled the air. Gravel rattled underneath my shoes as I hurried to the car. Before jumping behind the wheel, I noticed Jarkko's backpack on the roof of my car. So typical, I smiled to myself, as I carried the backpack back inside. When I left it on the floor in the hallway, I noticed Jarkko quickly stopping a phone call and blushing almost as if guilty. However, Jarkko assured that someone had just called the wrong number. I was in a hurry so I asked no further questions, but hasted to my car.

Engine ran steadily, as I drove through the peaceful rural landscape. The terrain varied with forests and fields. On ridges grew pines and in valleys dense spruce. In other places the road crossed over small rapids. Nature already started to turn green, much to the influence of the spring sun.

I reflected on the behaviour of Jarkko this morning: his sudden disconnection of the phone call and blushing as if guilty. I wonder whether Jarkko had something inappropriate going on with someone. Would he guess that he had awakened my doubts, and, if so, would he be scared enough to terminate the relationship. I wondered if I would dare to spy from Jarkko's phone who the caller was if I had the opportunity? Would Jarkko suspect that and empty his phone-call records? Maybe I should confront Jarkko if the call data had been cleared.

I woke to the reality from these gloomy reflections, when a large tree fell with a crash across the road. Brakes screeched as I struggled to stop the car before a crash! I climbed out of the car to see what had downed the tree. To my surprise, I saw a brown-haired, square teethed creature at the foot of the tree, and the root of the tree had bite marks. Beaver! When I approached the beaver, he fled deeper into the forest. Although we had already lived for about five years in Canada, this was the first time I saw a beaver. Shaking my head, I hastened back to the car, I turned the car around and I planned an alternate route to work in my mind. At the same time I took the phone from my pocket and I called the local emergency number, to declare fallen timber.

As I entered the elevator in the parking garage of my working place, I came across Mark, a handsome man with whom I had had a secret romance in the fall. Mark had begun to suspect that his wife knows something, and then we agreed that we take a break and let things cool down. I suspected, however, that Mark was just tired of me, and that his wife had actually not suspected anything. It may well be that Mark had another lover, and he did not want to mix things up too much. But I hid my doubts and I chit chatted with him. Mark and I were going to the same meeting. On the way to a meeting room, a chain-saw that someone had left in the corridor caught my attention.

## REFERENCES

1. Sumby, W.H. and Pollack, I. No Title. J. Acoust. Soc. Am. 26, 212–215 (1954).

2. Bernstein, L. E., Auer, E. T. & Takayanagi, S. Auditory speech detection in noise enhanced by lipreading. Speech Commun. 44, 5–18 (2004).

3. Ross, L. A. et al. Do You See What I Am Saying? Exploring Visual Enhancement of Speech Comprehension in Noisy Environments. (2007). doi:10.1093/cercor/bhl024

4. Strelnikov, K. et al. deafness after adult cochlear implantation. 3682–3695 (2013). doi:10.1093/brain/awt274

5. Bernstein, L. E. & Liebenthal, E. Neural pathways for visual speech perception. Front. Neurosci. 8, 1–18 (2014).

6. Files, B. T., Tjan, B. & Bernstein, L. E. Visual speech discrimination and identification of natural and synthetic stimuli. 6, 1–18 (2015).

7. Campbell, R. & Mohammed, T. Speechreading for information gathering: a survey of scientific source. 1–22 (2010).

8. Woodhouse, L., Hickson, L. & Dodd, B. Review Review of visual speech perception by hearing and hearing-impaired people□: clinical implications. (2009). doi:10.1080/13682820802090281

9. Ronnberg, J., Andersson, J., Samuelsson, S., Soderfeldt, B., Lyxell, B., & Risberg, J. A speechreading expert: The case of MM. J. Speech, Lang. Hear. Res. 42, (1999).

10. Altieri, N. a, Pisoni, D. B. & Townsend, J. T. Some normative data on lip-reading skills (L). J. Acoust. Soc. Am. 130, 1–4 (2011).

11. Anderson, C. A., Wiggins, I. M., Kitterick, P.T. & Hartley, D. E. H. Adaptive benefit of cross-modal plasticity following cochlear implantation in deaf adults. 114, (2017).

12. Lyxell, B. and Rönnberg, J. J. Information-processing skill and speechreading. Br. J. Audiol. 23, (1989).

13. Bernstein, L. E., Demorest, M. E. & Tucker, P. E. Speech perception without hearing. Percept. Psychophys. 62, 233–252 (2000).

14. Andersson, U. & Lidestam, B. Bottom-Up Driven Speechreading in a Speechreading Expert□: The Case of AA ( JK023 ). 214–224 (2005).

15. Kyle, F. E. & Harris, M. Predictors of reading development in deaf children: A 3-year longitudinal study. J. Exp. Child Psychol. 107, 229–243 (2010).

16. Feld, J.E. and Sommers, M. S. NIH Public Access. 4, 1555–1565 (2011).

17. Rönnberg, J. The Ease of Language Understanding ( ELU ) model: theoretical, empirical, and clinical advances. 7, 1–17 (2013).

18. Mohammed, T., Campbell, R., Macsweeney, M., Barry, F. & Coleman, M. Speechreading and its association with reading among deaf, hearing and dyslexic individuals. Clin. Linguist. Phon. 20, 621–630 (2006).

19. Campbell, R. Speechreading and the Bruce-Young model of face recognition: Early findings and recent developments. Br. J. Psychol. 102, 704–710 (2011).

20. Chu, Y. H. et al. Effective cerebral connectivity during silent speech reading revealed by functional magnetic resonance imaging. PLoS One 8, (2013).

21. Calvert, G.A., Bullmore E.T., Brammer, M.J, Campbell, R., Williams, S.C.R., McGuire, P.K, Woodruff, P.W.R., Iversen, S.D., David, A. S. Activation of Auditory Cortex During Silent Lipreading. Science (80-.). 276, 593.596 (1997).

22. Möttönen, R., Krause, C. M., Tiippana, K. & Sams, M. Processing of changes in visual speech in the human auditory cortex. Cogn. Brain Res. 13, 417–425 (2002).

23. Sams, M., Aulanko, R., Hämäläinen, M., Hari, R., Lounasmaa, O.V., Sing-Teh Lu, J. Seeing speech: visual information from lip movements modifies activity in the human auditory cortex. Neurosci. Lett. 10, 141.145 (1991).

24. Pekkola, J., Ojanen, V., Autti, T., Jääskeläinen, I.P., Möttönen, R., Tarkiainen, A., & Sams, M. Primary auditory cortex activation by visual speech: and fMRI study at 3T. Neuroreport 16, 125.128 (2005).

25. Paulesu, E. et al. A Functional-Anatomical Model for Lipreading. 2005–2013 (2003).

26. Nishitani, N. and Hari, R. R. Viewing Lip Forms□: Cortical Dynamics. Neuron 36, 1211–1220 (2002).

27. Watkins, K. E., Strafella, A. P. & Paus, T. Seeing and hearing speech excites the motor system involved in speech production. 41, 989–994 (2003).

28. Skipper, J. I., van Wassenhove, V., Nusbaum, H. C. & Small, S. L. Hearing lips and seeing voices: how cortical areas supporting speech production mediate audiovisual speech perception. Cereb. Cortex 17, 2387–2399 (2007).

29. Callan, D. E., Jones, J. A., Callan, A. & Nusbaum, H. C. Multisensory and modality specific processing of visual speech in different regions of the premotor cortex. 5, 1–10 (2014).

30. Calvert, G. a & Campbell, R. Reading speech from still and moving faces: the neural substrates of visible speech. J. Cogn. Neurosci. 15, 57–70 (2003).

31. Ludman, C. N. et al. Lip-reading ability and patterns of cortical activation studied using fMRI. Br. J. Audiol. 34, 225–30 (2000).

32. Hall, D. a, Fussell, C. & Summerfield, a Q. Reading fluent speech from talking faces: typical brain networks and individual differences. J. Cogn. Neurosci. 17, 939–953 (2005).

33. Friederici, A. D. The brain basis of language processing: From structure to function. Physiol. Rev. 91, 1357–92 (2011).

34. Poeppel, D., Emmorey, K., Hickok, G. & Pylkkanen, L. Towards a New Neurobiology of Language. J. Neurosci. 32, 14125–14131 (2012).

35. DeWitt, I. & Rauschecker, J. P. Phoneme and word recognition in the auditory ventral stream. Proc. Natl. Acad. Sci. 109, E505–E514 (2012).

36. Leonard, M. K. & Chang, E. F. NIH Public Access. 18, 472–479 (2015).

37. Mesgarani, N., Cheung, C., Johnson, K., Chang, E. F. & Francisco, S. HHS Public Access. 343, 1006–1010 (2015).

38. Overath, T., Mcdermott, J. H., Zarate, J. M. & Poeppel, D. HHS Public Access. 18, 903–911 (2016).

39. Capek, C. M. et al. Cortical circuits for silent speechreading in deaf and hearing people. Neuropsychologia 46, 1233–1241 (2008).

40. Hasson, U., Malach, R. & Heeger, D. J. Reliability of cortical activity during natural stimulation. Trends Cogn. Sci. 14, 40–48 (2010).

41. Hasson, U., Nir, Y., Levy, I., Fuhrmann, G. & Malach, R. Natural Vision.

42. Kauppi, J.-P., Jääskeläinen, I. P., Sams, M. & Tohka, J. Inter-subject correlation of brain hemodynamic responses during watching a movie: localization in space and frequency. Front. Neuroinform. 4, 5 (2010).

43. Pajula, J., Kauppi, J. P. & Tohka, J. Inter-subject correlation in fMRI: Method validation against stimulus-model based analysis. PLoS One 7, (2012).

44. Price, C. J. The anatomy of language: A review of 100 fMRI studies published in 2009. Ann. N. Y. Acad. Sci. 1191, 62–88 (2010).

45. Yousry, T.A., Schmid, U.D., Alkadhi, H., Schmidt, D., Peraud, A., Buettner, A., & Winkler, P. Localization of the motor hand area to a know on the precentral gyrus. A new landmark. Brain 120, 141–57 (1997).

46. Fox, P. T. et al. Location-probability profiles for the mouth region of human primary motor-sensory cortex: model and validation. Neuroimage 13, 196–209 (2001).

47. Hickok, G. & Poeppel, D. The cortical organization of speech processing. Nat. Rev. Neurosci. 8, 393–402 (2007).

48. Binder, J. R., Desai, R. H., Graves, W. W. & Conant, L. Where Is the Semantic System□? A Critical Review and Meta-Analysis of 120 Functional Neuroimaging Studies. (2009). doi:10.1093/cercor/bhp055

49. Rodd, J. M., Vitello, S., Woollams, A. M. & Adank, P. Localising semantic and syntactic processing in spoken and written language comprehension: An Activation Likelihood Estimation meta-analysis. Brain Lang. 141, 89–102 (2015).

50. Snijders, T. M. et al. Retrieval and unification of syntactic structure in sentence comprehension: An fMRI study using word-category ambiguity. Cereb. Cortex 19, 1493–1503 (2009).

51. Wilson, S. M., Molnar-Szakacs, I. & Iacoboni, M. Beyond superior temporal cortex: intersubject correlations in narrative speech comprehension. Cereb. Cortex 18, 230–242 (2008).

52. Honey, C. J., Thompson, C. R., Lerner, Y. & Hasson, U. Not Lost in Translation: Neural Responses Shared Across Languages. J. Neurosci. 32, 15277–15283 (2012).

53. Regev, M., Honey, C. J., Simony, E. & Hasson, U. Selective and invariant neural responses to spoken and written narratives. J. Neurosci. 33, 15978–88 (2013).

54. Huth, A. G., Heer, W. A. De, Griffiths, T. L., Theunissen, F. E. & Jack, L. Natural speech reveals the semantic maps that tile human cerebral cortex. Nature 532, 453–458 (2016).

55. Campbell, R. Everyday speechreading: understanding seen speech in action. Scand. J. Psychol. 39, 163–167 (1998).

56. Poeppel, D. The analysis of speech in different temporal integration windows: Cerebral lateralization as ‘asymmetric sampling in time’. Speech Commun. 41, 245–255 (2003).

57. Weikum, W. M. et al. Visual Language Discrimination in Infancy -- Weikum et al. 316 (5828)_ 1159 -- Science. Science (80-). 25, (2007).

58. Munhall, K. G., Jones, J. a, Callan, D. E., Kuratate, T. & Vatikiotis-Bateson, E. Visual prosody and speech intelligibility: head movement improves auditory speech perception. Psychol. Sci. a J. Am. Psychol. Soc. /APS 15, 133–137 (2004).

59. Scott, S. K., Blank, C. C., Rosen, S. & Wise, R. J. S. Identification of a pathway for intelligible speech in the left temporal lobe. 2400–2406 (2000).

60. Binder, J.R., Frost, J.A., Hammeke, T.A., Bellgovan, P.S.F., Springer, J. A., Kaufman, J. N. & Possing, E. T. Human Temporal Lobe Activation by Speech and Nonspeech Sounds. Cereb. Cortex 10, 512–528 (2000).

61. Kyle, F. E., Campbell, R., Mohammed, T., Coleman, M. & Ad Macsweeney, M. Speechreading Development in Deaf and Hearing Children: Introducing the Test of Child Speechreading. J. Speech Hear. Res. 56, 416–426 (2013).

62. Lu, K.-H., Hung, S.-C., Wen, H., Marussich, L. & Liu, Z. Influences of High-Level Features, Gaze, and Scene Transitions on the Reliability of BOLD Responses to Natural Movie Stimuli. PLoS One 11, e0161797 (2016).

63. O’Sullivan, A. E., Crosse, M. J., Di Liberto, G. M. & Lalor, E. C. Visual Cortical Entrainment to Motion and Categorical Speech Features during Silent Lipreading. Front. Hum. Neurosci. 10, 679 (2017).

64. Venezia, J. H. et al. Network for Sensorimotor Integration of Visual Speech. 196–207 (2017). doi:10.1016/j.neuroimage.2015.11.038.Perception

65. Skipper, J. I., Nusbaum, H. C. & Small, S. L. Listening to talking faces: motor cortical activation during speech perception. Neuroimage 25, 76–89 (2005).

66. Käuramaki, J. et al. Lipreading and covert speech production similarly modulate human auditory-cortex responses to pure tones. J. Neurosci. 30, 1314–1321 (2010).

67. Booth, J. R., Wood, L., Lu, D., Houk, J. C. & Bitan, T. processing. 1133, 136–144 (2008).

68. Fedorenko, E., Behr, M. K. & Kanwisher, N. Functional specificity for high-level linguistic processing in the human brain. Proc. Natl. Acad. Sci. U. S. A. 108, 16428–16433 (2011).

69. Fairhall, S. L. & Caramazza, A. Brain Regions That Represent Amodal Conceptual Knowledge. 33, 10552–10558 (2013).

70. Mahowald, K. & Fedorenko, E. Reliable individual-level neural markers of high-level language processing: A necessary precursor for relating neural variability to behavioral and genetic variability. Neuroimage 139, 74–93 (2016).

71. Binder, J. R. & Desai, R. H. The neurobiology of semantic memory. Trends Cogn. Sci. 15, 527–536 (2011).

72. Regev, M., Honey, C. J., Simony, E. & Hasson, U. Selective and invariant neural responses to spoken and written narratives. J. Neurosci. 33, 15978–88 (2013).

73. Raichle, M. E. et al. A default mode of brain function. Proc. Natl. Acad. Sci. U. S. A. 98, 676–82 (2001).

74. Utevsky, A. V, Smith, D. V & Huettel, S. A. Precuneus Is a Functional Core of the Default-Mode Network. 34, 932–940 (2014).

75. Simony, E. et al. network during narrative comprehension. Nat. Commun. 7, 1–13 (2016).

76. Yeshurun, Y. et al. Same Story, Different Story. Psychol. Sci. 95679761668202 (2017). doi:10.1177/0956797616682029

77. Buckner, R. L. The cerebellum and cognitive function: 25 years of insight from anatomy and neuroimaging. Neuron 80, 807–815 (2013).

78. Stoodley, C. J., Valera, E. M. & Schmahmann, J. D. NeuroImage Functional topography of the cerebellum for motor and cognitive tasks□: An fMRI study. Neuroimage 59, 1560–1570 (2012).

79. Wang, D., Buckner, R. L. & Liu, H. Cerebellar asymmetry and its relation to cerebral asymmetry estimated by intrinsic functional connectivity. J. Neurophysiol. 109, 46–57 (2013).

80. Kotz, S. A., Stockert, A., Schwartze, M. & Kotz, S. A. Cerebellum, temporal predictability and the updating of a mental model. (2014).

81. Nummenmaa, L. et al. Emotional speech synchronizes brains across listeners and engages large-scale dynamic brain networks. Neuroimage 102, 498–509 (2014).

82. Boldt, R. et al. Listening to an Audio Drama Activates Two Processing Networks, One for All Sounds, Another Exclusively for Speech. PLoS One 8, 1–10 (2013).

83. Bar, M. The proactive brain: using analogies and associations to generate predictions. Trends Cogn. Sci. 11, 280–289 (2007).

84. Oldfield, R. C. The assessment and analysis of handedness: the Edinburgh inventory. Neuropsychologia 9, 97–113 (1971).

85. Sims, D. The validation of the CID everyday sentence test for use with the severely hearing impaired. 70–79 (1975).

86. Lonka, E. Hard-of-hearing adult and learning to speechread. (1993).

87. Lahnakoski, J. M. et al. Naturalistic fMRI Mapping Reveals Superior Temporal Sulcus as the Hub for the Distributed Brain Network for Social Perception. Front. Hum. Neurosci. 6, 1–14 (2012).

88. Power, J. D. et al. NeuroImage Methods to detect, characterize, and remove motion artifact in resting state fMRI. Neuroimage 84, 320–341 (2014).

89. Eklund, A., Nichols, T. E. & Knutsson, H. Cluster failure: Why fMRI inferences for spatial extent have inflated false-positive rates. Proc. Natl. Acad. Sci. 113, 201602413 (2016).

90. Mantel, N. The detection of disease clustering and a generalized regression approach. Cancer Res. 27, 209–20 (1967).

